# In search for multifunctional lncRNAs

**DOI:** 10.1101/2024.07.11.603032

**Authors:** Bharat Ravi Iyengar

## Abstract

Long non-coding RNAs (lncRNAs) were so named because at the time of their discovery, no corresponding protein products were known. Despite the lack of evidence for translation, many lncRNAs perform essential cellular functions such as regulation of gene expression. Recent studies show that many lncRNAs, including those with known regulatory functions, bind to ribosomes and are translated, suggesting that RNAs can perform two different kinds of functions (a phenomenon known in proteins, called moonlighting). Using a formal mathematical model, I show that execution of one function limits that of the other. However, an RNA can transition from one function to the other, simply by a spatiotemporal regulation of its interacting partners. I further studied the properties of proteins encoded in functional human lncRNAs and found that many of them have complex sequences, and some of them can even adopt stable 3D structures. These findings may encourage further exploration of moonlighting lncRNAs, their regulation, and their role in the evolution of new protein coding genes.

## Main

One of the best known functions of ribonucleic acid (RNA) is to serve as a template for protein synthesis (mRNA). The RNAs that are not known to encode a protein are designated as non-coding RNAs (ncRNAs, Mattick and Makunin, 2006). Encoding a protein is not the only function an RNA can perform. The best example would be that of ribosomal RNAs (rRNAs) and transfer RNAs (tRNAs) that are pivotal for protein synthesis. Generally, the functions of non-coding RNAs are diverse, ranging from essential housekeeping functions such as splicing (snRNAs), ribosome biogenesis (snoRNAs), and translation (rRNA, tRNA), to specific functions such as regulation of gene expression (miRNAs). Long non-coding RNAs (lncRNAs) are ncRNAs longer than 200nt, several of which are involved in gene regulation (lncRNAs, Statello *et al*., 2020; Mattick *et al*., 2023). It has been shown that many lncRNAs bind to ribosomes and are translated to synthesize proteins (Ruiz-Orera *et al*., 2014; Ingolia *et al*., 2014; Patraquim *et al*., 2022). Thus their classification as non-coding RNAs may be inaccurate, and may need reassessment (Jalali *et al*., 2016; Patraquim *et al*., 2022). Interestingly, among these ribosome-associated (and translated) lncRNAs are several lncRNAs that perform other distinct functions. For example, a ribosome associated lncRNA, *bxd* is involved in repressing the transcription of the developmental gene Ultrabithorax (*Ubx*) in *Drosophila melanogaster* (Patraquim *et al*., 2022). Similarly, the mammalian lncRNA *Malat1*, that is known to regulate gene expression and cell cycle, can also associate with ribosomes and synthesize a protein (Ingolia *et al*., 2014). It is yet to be determined if the proteins produced from the translation of different lncRNAs have any role in cellular physiology or if they are simply accidental by-products. Nonetheless, it is clear that an RNA molecule can perform two distinct functions. Such a non-singular functionality, called “moonlighting”, is well known in proteins (Jeffery, 2017). An important question about moonlighting is whether the two functions of a biomolecule can be simultaneously executed by the same molecule. Specifically in case of lncRNAs, it is important to understand if engagement of an lncRNA in protein translation, would compromise its other cellular function (for example, gene regulation). Many biomolecules interact with specific partner molecules to perform a function. For example, the lncRNA *Chaserr* binds to the chromatin remodelling protein *Chd2* (Rom *et al*., 2019). One can argue that binding to a ribosome may prevent an lncRNA to associate with its other binding partners, and its localization to specific cellular compartments where it performs its functions. Conversely, an lncRNA’s cognate function and binding partners, may prevent it from binding to the ribosome and being translated to a protein.

I developed a a simple model of competition to explain the conflict between the two possible functions of an lncRNA. In this model, a hypothetical RNA (*R*) binds to an RNA binding protein (RBP, denoted by *X*) to form a ribonucleoprotein complex (*C*). The lncRNA can also bind to a ribosome to form a productive translation elongation complex (*E*), that translates an open reading frame (ORF) in the RNA. The ribosome disengages from the RNA when the elongation complex reaches a stop codon, while releasing a molecule of the translated protein. Because the free ribosome is usually in surplus, its concentration would not play a major role in the dynamics of lncRNA binding. This model of lncRNA partitioning illustrated in Figure 1A, can be described by the following set of ordinary differential equations:

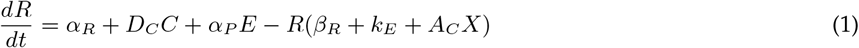

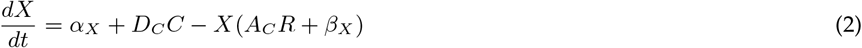

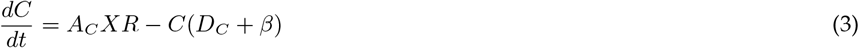

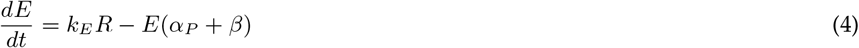

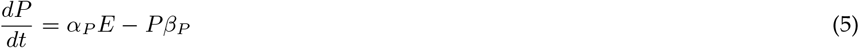

**Figure 1:**
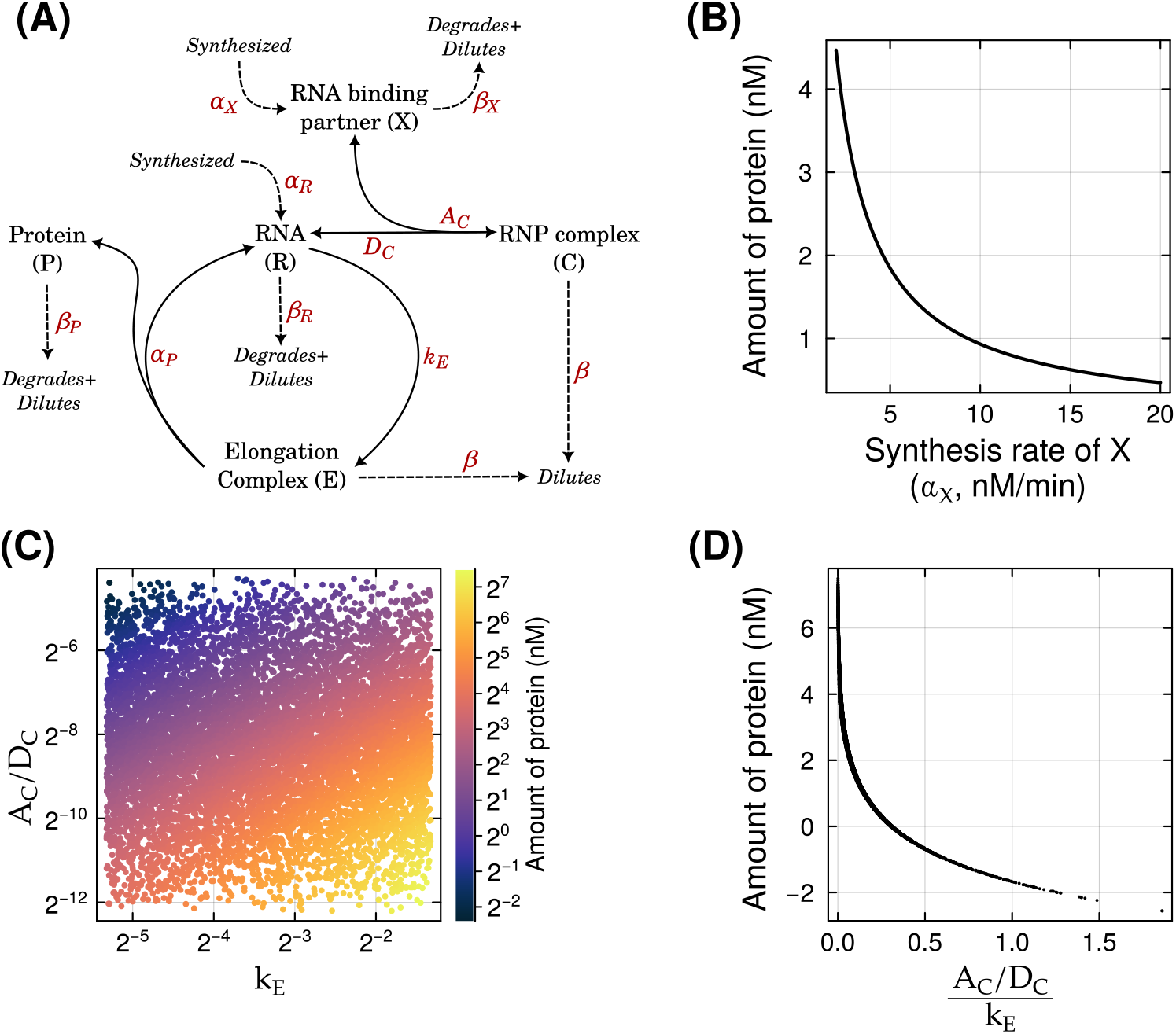
Model of competition between RNA binding protein and ribosome. **(A)** Illustration of the competition model, that is formally described in Equations 1 – 5. The parameters that denote the various biochemical reactions are described in Table 1. Here, dilution refers to dilution of biomolecules due to cell division. The RNP (*X*) and the ribosome compete for binding the RNA. **(B)** A higher rate of RNP expression (*α*_*X*_, horizontal axis) causes lower translation of the protein *P* (vertical axis).**(C)** Stronger binding to RNP (*A*_*C*_*/D*_*C*_ = *K*_*AC*_, vertical axis, log_2_-scaled), and slower initiation of translation (*k*_*E*_, horizontal axis, log_2_-scaled), leads to lower translation of protein *P* (color gradient, log_2_-scaled), and *vice versa*. The scatter plot displays 10000 data points that describe the steady state solutions with 10000 parameter sets, where we randomly sampled the parameters *A*_*C*_, *D*_*C*_ and *k*_*E*_ from a multivariate log-uniform distribution ranging from 1/4 to 4 times their default values. We kept the other parameters at their default values. **(D)** is a summary of panel **(B)**. Here, the horizontal axis (log_2_-scaled) shows the amount of protein and the vertical axis shows the *K*_*AC*_*/k*_*E*_ ratio.

Solutions of these equations at steady state (equating each equation to zero), describe the long term behavior of the system, and are dependent on the values of the different parameters (Table 1). The competition between the ribosome and the RBP can be described by four parameters. The first parameter describes the rate at which the RBP (*X*) is synthesized (*α*_*X*_) that determines its total amount in the cell. The second parameter (*A*_*C*_) describes the rate at which RNA binds to the RBP to form the complex (*C*). The third parameter (*D*_*C*_) denotes the rate at which the complex dissociates into its two individual constituents. The fourth parameter (*k*_*E*_) describes the rate at which translation of an ORF in the RNA is initiated.

**Table 1:** Values of different parameters (rate constants) used in the Equations 1 – 5. These values are estimated from published data in the corresponding references, and are biologically reasonable. The standard abbreviations, nM and min, stand for nanomolars and minutes, respectively.

| Parameter (rate constant) | Symbol | Default value | Estimated from |
| --- | --- | --- | --- |
| RNA length | - | 1000 nt | assumption: variable |
| ORF length | - | 100 codons | assumption: variable |
| ORF position | - | 100 nt | assumption: variable |
| RNA (R) synthesis | $\alpha_R$ | $\frac{2.5}{\text{RNA length}} \text{ nM} \cdot \text{min}^{-1}$ | Veloso et al. (2014) |
| Synthesis of RBP (X) | $\alpha_X$ | $4.78 \text{ nM} \cdot \text{min}^{-1}$ | assumption: variable |
| Translation of Protein (P) from RNA | $\alpha_P$ | $\frac{300}{\text{ORF length}} \text{ min}^{-1}$ | Gerashchenko et al. (2020) |
| Translation initiation | $k_E$ | $\frac{60}{6 \times \text{ORF position}} \text{ min}^{-1}$ | Vassilenko et al. (2011) |
| Complex formation | $A_C$ | $10^{-3} \text{ nM}^{-1} \cdot \text{min}^{-1}$ | Jain et al. (2017) |
| Complex dissociation | $D_C$ | $0.3 \text{ min}^{-1}$ | Jain et al. (2017) |
| Degradation/dilution of RNA | $\beta_R$ | $0.0017 \text{ min}^{-1}$ | division time of 10h |
| Degradation/dilution of Protein (P) | $\beta_P$ | $0.0017 \text{ min}^{-1}$ | division time of 10h |
| Degradation/dilution of RBP | $\beta_X$ | $0.0017 \text{ min}^{-1}$ | division time of 10h |
| General dilution | $\beta$ | $0.0017 \text{ min}^{-1}$ | division time of 10h |

A high expression of the RBP (*X*) leads to a smaller amount of translated protein (*P*), because more RNP is available to bind to the RNA, and prevent it from being translated (Figure 1B). A higher expression of the RBP will cause lower translation of the RNA as long as all the RBP molecules are not already bound by the RNA, or if there are no regulatory mechanisms such as self-inhibition. More specifically, the rate of RNA-RBP binding depends on the amount of free RBP. A stable ribonucleoprotein complex can form due to a fast association of the constituents and a slow dissociation of the complex. Specifically, the ratio of the association and dissociation rate constants (*A*_*C*_*/D*_*C*_), that is the equilibrium association constant (*K*_*AC*_), is a good indicator of the complex’s stability. A more stable complex would sequester the RNA better, which in turn would prevent the translation of the RNA. The rate of translation initiation *k*_*E*_ determines how quickly the RNA commits to translation, from which it would be released only when the entire ORF is translated. The competition between RBP and ribosome can thus be described by the value of *K*_*AC*_ relative to *k*_*E*_. Specifically, the amount of translated protein is minimum when the *K*_*AC*_ is high and *k*_*E*_ is low (Figure 1C). Conversely, the protein’s expression is highest when *K*_*AC*_ is low and *k*_*E*_ is high. More specifically, the amount of translated protein decreases super-exponentially with increasing values of *K*_*AC*_*/k*_*E*_ ratio (Figure 1D). While the association and dissociation rate constants completely depend on the biochemical properties of the RBP and the RNA (including their 3D structures), the rate of translation initiation can depend on the location of the ORF in the RNA. This assumption follows the widely accepted model of eukaryotic translation initiation, where the ribosome binds to a capped RNA and slides along its length (at an approximate rate of 6-8nt per second) until it encounters a start codon (Vassilenko *et al*., 2011). Thus mRNAs, that are optimized for being translated would contain the ORF close to their 5’ end. Conversely, an lncRNA would lack such an optimal placement of the ORF. Analyses of translated lncRNAs in Drosophila show that highly efficiently translated ORFs are indeed located close to the 5’ end of the RNA (Patraquim *et al*., 2022). Nucleotide composition around the start codon also has an influence on translation initiation efficiency. Specifically, a sequence of seven nucleotides including the start codon itself, known as Kozak consensus sequence, determine translation initiation efficiency (Kozak, 1986; Noderer *et al*., 2014; Acevedo *et al*., 2018). In an RNA with many ORFs, the Kozak sequence determines which one of them is most efficiently translated. To understand if mRNAs are more optimized for translation than lncRNAs, I analysed the position of ORFs in mRNAs and lncRNAs, and the strength of the Kozak sequences (KCS) flanking their start codons. Specifically, I estimated the correlation between KCS (Noderer *et al*., 2014; Acevedo *et al*., 2018) and ORF position in the annotated mRNAs from *Drosophila melanogaster* (FlyBase, Gramates *et al*., 2022) and *Homo sapiens* (RefSeq, O’Leary *et al*., 2015). I found that Drosophila ORFs with a large KCS are frequently present towards the 5’ end of the mRNAs (Spearman *ρ* = *−*0.16, *P <* 10^*−*15^; Figure 2A). This correlation was weaker for human mRNAs but was nonetheless statistically significant (Spearman *ρ* = *−*0.043, *P <* 10^*−*15^). Next, I analysed the correlation between ORF position and KCS for the set of all the annotated lncRNAs in Drosophila (FlyBase, Gramates *et al*., 2022) and found that ORFs with a large KCS were usually located towards the 3’ end of the RNA (Spearman *ρ* = 0.0383 *P* = 5.7 *×* 10^*−*5^; Figure 2B). This was not the case for the annotated human non-coding RNAs in RefSeq, where the ORF position was significantly negatively correlated with the KCS (Spearman *ρ* = *−*0.037, *P <* 10^*−*15^). However, when I restricted this analysis to a set of experimentally validated functional human lncRNAs (lncRNAWiki 2.0, Liu *et al*., 2021), the correlation was no longer significant (Spearman *ρ* = *−*0.003, *P* = 0.56). Thus it appears that ORFs encoded in functional lncRNAs are indeed not optimized for translation, probably because their translation can possibly interfere with the regulatory functions of the RNA. This leads back to the original question of whether an RNA can perform both the translational and non-translational functions efficiently. The model and the data suggest that this is not very likely. However, it also shows that by tuning the expression of the RBP (or any other binding partner), a cell can switch a translating RNA to a non-translating RNA (Figure 1A). Such a switching behavior has indeed been shown to exist in Drosophila where the some lncRNAs are translated only in specific developmental stages (Patraquim *et al*., 2022). LncRNAs can also “moonlight” in other ways. For example, they can associate with different binding partners under different conditions, and can thus perform different functions (Cheng and Leung, 2018; Balcerak *et al*., 2019). In these cases too, the competition between two binding partners will impose a trade-off between the two functions, A spatio-temporal segregation of the two functions, by regulated expression of the binding partners can resolve this trade-off, which can be resolved by regulation of the binding partners.

**Figure 2:**
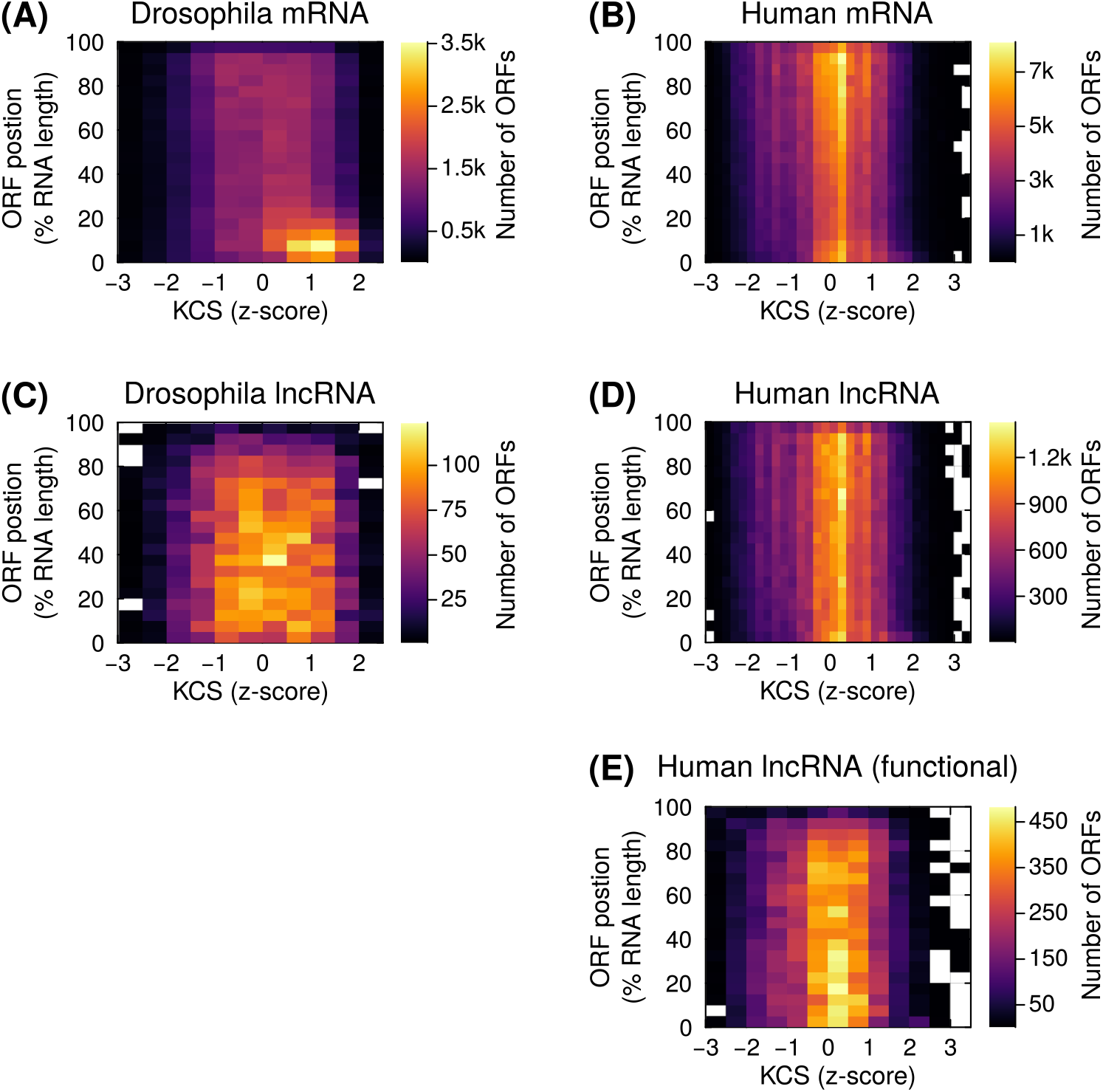
ORFs with high KCS are generally not optimally placed. Panels **A** – **E** denote ORFs in Drosophila mRNAs (FlyBase), human mRNAs (RefSeq), Drosophila lncRNAs (FlyBase), human lncRNAs (RefSeq) and functional human lncRNAs (lncRNAWiki), respectively. In each panel, the horizontal axis denotes the strength of the Kozak consensus sequence (normalized as z-score), the vertical axis denotes the start of the ORF relative to the length of the parent RNA, and the color scale denotes the number of ORFs (yellow = many, black = few).

To understand if functional lncRNAs can also encode functional proteins, I computationally analysed the properties of protein sequences encoded in a set of experimentally validated functional human lncRNAs (lncRNAWiki 2.0, Liu *et al*., 2021). I restricted this analysis to 7669 lncRNAs whose interaction target was identified as a protein coding gene, an RNA or a protein. Henceforth, all my references to lncRNA encoded proteins describe the proteins encoded in these 7669 lncRNAs. First, I analysed the lengths of these lncRNA encoded proteins, and found that they are significantly shorter (median 47 amino acids) than conserved human proteins in RefSeq database (median 514 amino acids; one sided Mann-Whitney U test, *P <* 10^*−*15^; Figure 3A). The frequency of lncRNA encoded proteins exponentially reduced with their lengths as expected (Iyengar and Bornberg-Bauer, 2023; Lebherz *et al*., 2024).

**Figure 3:**
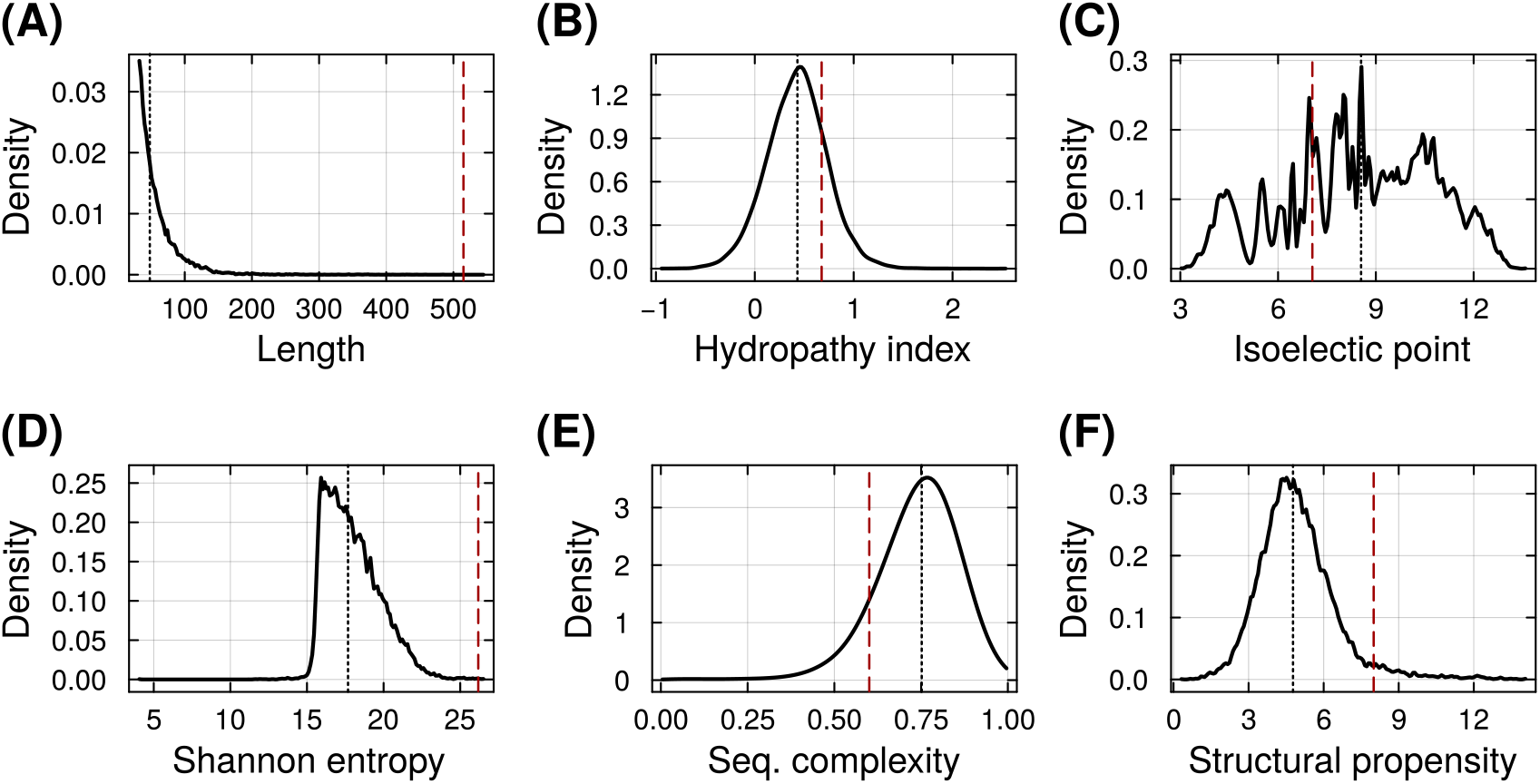
Properties of proteins encoded in a set of 7669 functional human lncRNAs denoted as density plots – **(A)** length, **(B)** hydropathy index, **(C)** isoelectric point, **(D)** Shannon entropy, **(E)** linguistic complexity, and **(F)** structure propensity. The black dotted vertical line in each panel denotes the median value of the corresponding property among the lncRNA encoded proteins. The dark red dashed vertical line in panels **A** – **E** denotes the median value of the corresponding properties among human RefSeq proteins, whereas in panel **F**, it denotes the cutoff of 8 that delineates structured and disordered proteins (Redl *et al*., 2023).

**Figure 4:**
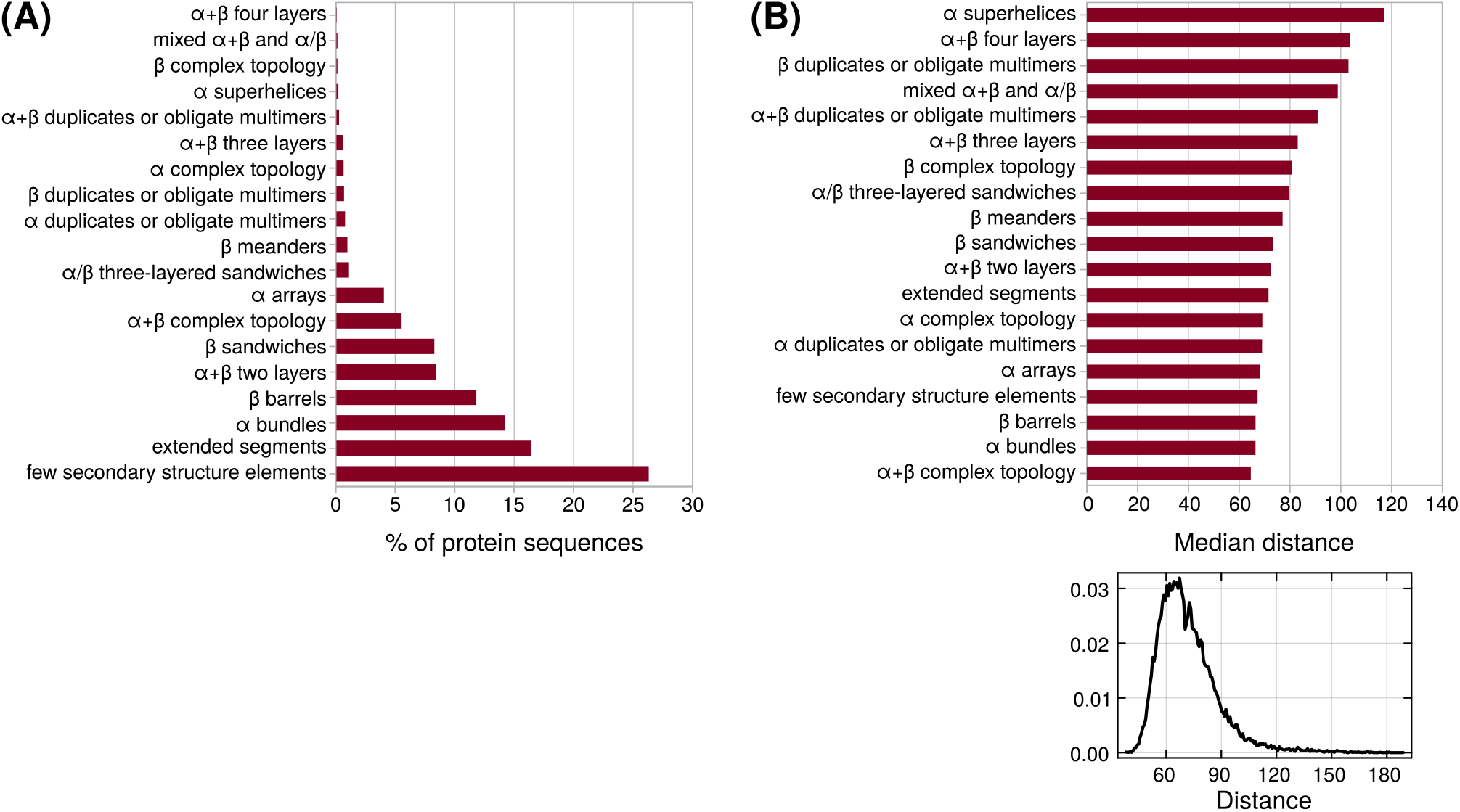
ECOD domains nearest to lncRNA encoded proteins, identified using the ESM2 protein language model. **(B)** Percentage of proteins (horizontal axis) in each ECOD domain class (vertical axis). **(C)** Median Manhattan (L1) distance between proteins and their nearest ECOD domains (horizontal axis) from each domain class (vertical axis). The plot below shows distribution of the distance of all the proteins to their nearest domains.

Random proteins are expected to have approximately 40% hydrophobic amino acids (Hochberg *et al*., 2020; Iyengar and Bornberg-Bauer, 2023), and are moderately hydrophilic (hydropathy index of *∼*0.38 given the locus GC content of 42%; Iyengar and Bornberg-Bauer, 2023). To test if this is also the case for lncRNA encoded proteins, I calculated the average hydropathy index for each protein (Wimley *et al*., 1996, octanol scale). I found that the hydropathy index of lncRNA encoded proteins (median 0.43) was significantly smaller than that of Ref-Seq proteins (median 0.68; one sided Mann-Whitney U test, *P <* 10^*−*15^; Figure 3B) but was comparable to the expected hydropathy index of random proteins (0.44) given the mean GC content of the lncRNAs (46%). Thus lncRNA encoded proteins tend to be more hydrophobic than both random proteins and conserved proteins. Next, I analysed the overall electrostatic charge distribution of lncRNA encoded proteins by calculating their isoelectric point. The predicted isoelectric point of lncRNA encoded proteins was significantly smaller than that of conserved proteins suggesting that the former are generally more acidic than the latter (one sided Mann-Whitney U test, *P <* 10^*−*15^; Figure 3C).

The amount of “information” contained in a protein sequence can be analysed from its composition. Proteins with low information (or complexity) contain repetitive sequences. Conversely, proteins with higher information/complexity contain a higher number of unique oligopeptide sequences (words). To quantify the amount sequence information, I used two metrics – Shannon entropy, and linguistic complexity. I found that the Shannon entropy of lncRNA encoded proteins was significantly smaller (median 17.7) than that of conserved proteins (median 26.2; one sided Mann-Whitney U test, *P <* 10^*−*15^; Figure 3D). However, the linguistic complexity of lncRNA encoded proteins was significantly larger (median 0.75) than that of conserved proteins (median 0.6; one sided Mann-Whitney U test, *P <* 10^*−*15^; Figure 3E). This discrepancy between the two metrics can be explained by the difference between the lengths of the two sets of proteins. While both metrics depend on the number of unique words in the sequence, Shannon entropy considers all the unique words that actually exist in a sequence, and is therefore, dependent on the sequence length. In contrast, linguistic complexity considers the number of unique words of a certain length that occur in a sequence, out of the total number of identically sized words that can exist in a sequence. To provide an analogy, a small book may contain less total information than a large book but the former can still be rich in vocabulary given its size. Analogously despite their small size, the short lncRNA encoded proteins are less repetitive than the larger conserved proteins.

It is widely believed that low complexity protein sequences tend to not adopt stable structures, because such sequences are uncommon among the proteins whose structures have been experimentally determined (Romero *et al*., 2000; Gonçalves-Kulik *et al*., 2022). While low sequence complexity may be sufficient to cause structural disorder, it is not necessary. Conversely, long repetitive sequences can form ordered structures (Mier *et al*., 2019). To understand if lncRNA encoded proteins are generally disordered or structured, I used a disorder prediction tool that is based on a protein language model (ADOPT, Redl *et al*., 2023). Using ADOPT, I found that most lncRNA encoded proteins are disordered (value below 8, Figure 3F) despite their high linguistic complexity. Although most lncRNA encoded proteins were disordered, some of them may evolve into structured proteins. To test this possibility, I mapped the lncRNA encoded proteins in the “sequence space” and found the nearest protein domain. Specifically, I used the ESM2 language model (Rice *et al*., 2000) to convert each protein sequence of any length into a numerical vector of a fixed length. These numerical values, called token embeddings, are obtained from the language model trained on protein sequences with known structures. I obtained the embeddings vector for lncRNA encoded proteins and the sequences of protein domains in the ECOD database (Cheng *et al*., 2014) that are 10 – 120 amino acids in length (thus excluding large domains that are unlikely to be found in short protein sequences). Next, I calculated Manhattan distance (or L1-norm) between the embedding vectors for each pair of lncRNA encoded protein and ECOD domain. Next, I identified the structural class of the ECOD domain that is nearest to any given lncRNA encoded protein. With this analysis, I found that 43% of lncRNA encoded proteins did not map to any structured domain. However, 14% and 11% of sequences mapped to α-bundles and β-barrels, respectively. The lncRNA encoded sequences were also generally closer in sequence space to these two domain classes than the other domains (except α+β complex topology). Overall, it appears that some lncRNA encoded proteins can adopt structures or can evolve to be structured. To further understand if some lncRNA encoded proteins are indeed structured, I predicted the structure of a few sequences using ESMfold (Lin *et al*., 2023). Specifically, I handpicked these proteins based on their low predicted disorder, and a relatively high KCS (with a few exceptions, Table 2). ESMfold predicted a high-confidence structure for most of these proteins, suggesting that they could be functional (Figure 5). The predicted structures also agreed well with the ECOD domain class descriptions (Table 2), suggesting that protein language models can be used for fast annotation of unknown protein sequences with domain architectures that they have or can evolve. An interesting case is that of protein encoded in the first ORF of the lncRNA CDKN2B-AS1. It has a ubiquitin like structure and can possibly be ligated to proteins via a mechanism similar to that of ubiquitin and other ubiquitin like proteins (such as SUMO, UFM and NEDD). The lncRNA CDKN2B-AS1 functions as an epigenetic regulator by interacting with the polycomb repressive complex, and is implicated in cancer progression (Hjazi *et al*., 2023).

**Table 2:** Description and some predicted properties of the lncRNA encoded proteins whose predicted structures are displayed in Figure 5.

| Figure 5 panel | lncRNA name | ENCODE id | ORF# | Structure propensity | KCS z-score | Nearest ECOD class | Nearest ECOD domain type | Distance |
| --- | --- | --- | --- | --- | --- | --- | --- | --- |
| A | HLA-F-AS1 | ENST00000414808 | 2 | 11.17 | -0.60 | $\alpha+\beta$ two layers | – | 84.38 |
| B | CDKN2B-AS1 | ENST00000651588 | 1 | 13.12 | 1.00 | $\alpha+\beta$ two layers | β-grasp | 43.82 |
| C | DUXAP8 | ENST00000456786 | 2 | 10.79 | 0.27 | $\alpha$ arrays | helix turn helix | 76.16 |
| D | FGD5-AS1 | ENST00000417835 | 1 | 9.63 | 0.59 | $\alpha$ arrays | helix turn helix | 169.77 |
| E | SBF2-AS1 | ENST00000663578 | 6 | 13.42 | 1.77 | few ss | RING/ U-box like | 136.81 |
| F | IFNG-AS1 | ENST00000674322 | 6 | 1.83 | 0.04 | $\alpha+\beta$ two layers | α-β plaits | 95.99 |
| G | SNHG14 | ENST00000654984 | 3 | 13.04 | 1.73 | $\alpha$ arrays | helix turn helix | 135.03 |
| H | LINC00662 | ENST00000658434 | 7 | 13.68 | 1.22 | mixed $\alpha+\beta$ and $\alpha/\beta$ | ribonuclease H like | 111.64 |
| I | MIR99AHG | ENST00000698124 | 10 | 14.07 | 0.31 | $\alpha$ arrays | helix turn helix | 112.46 |

**Figure 5:**
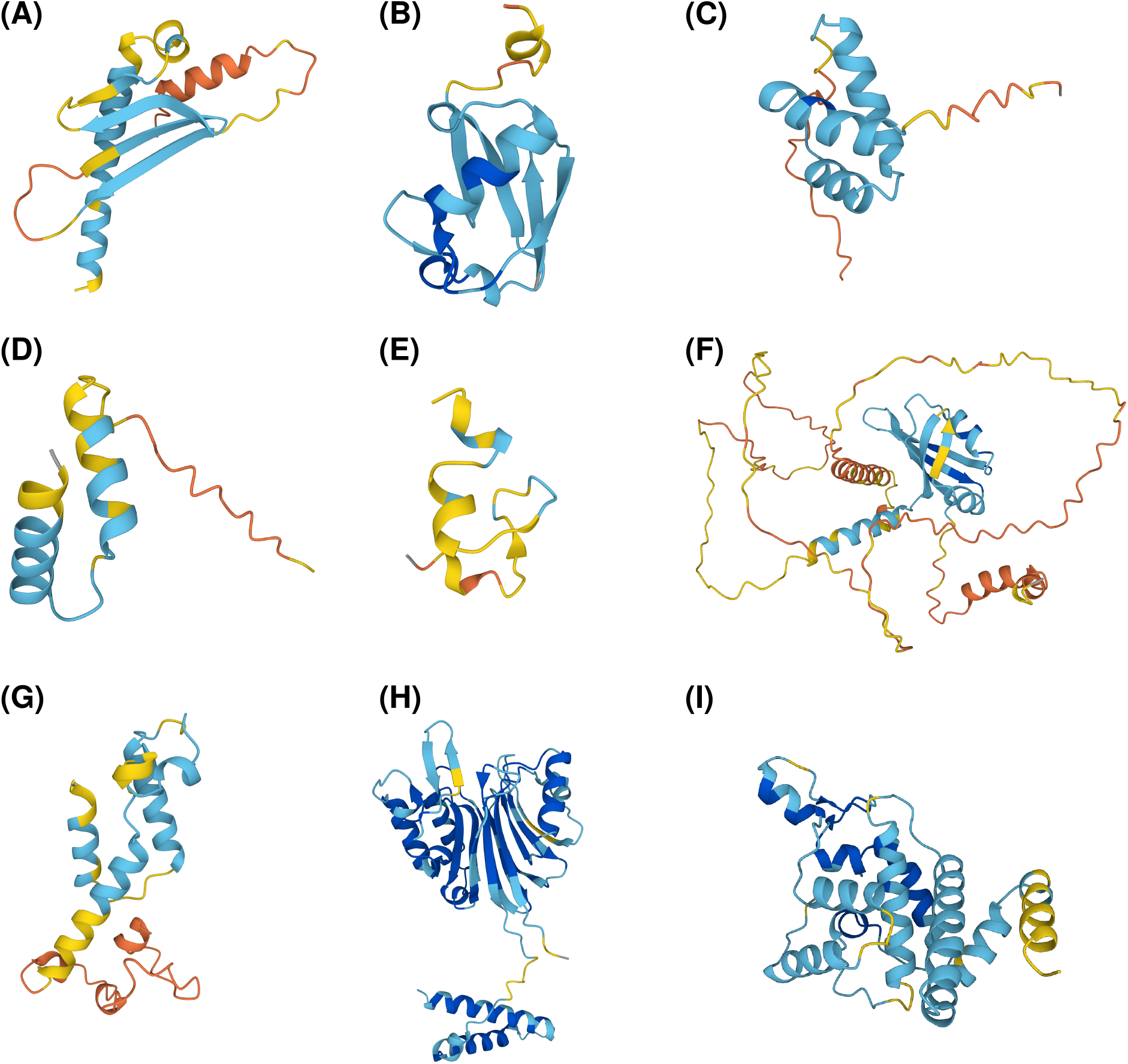
ESMfold predicted structures of a few selected lncRNA encoded proteins (Table 2). Secondary structures are depicted in the standard ribbon style. Blue and red shades denote high and low confidence (pLDDT) of the predicted structure, respectively.

Such moonlighting lncRNAs would be particularly interesting on the topic of *de novo* gene emergence (Van Oss and Carvunis, 2019; Schmitz and Bornberg-Bauer, 2017). In this phenomena, new protein coding genes (proto-genes) emerge from non-genic DNA sequences, by evolving features essential for transcription and translation. While transcriptional features are likely to emerge first (Iyengar and Bornberg-Bauer, 2023), they may also erode before translational features emerge. In functional lncRNAs, transcriptional features would be preserved via selection. Therefore emergence of proto-genes in lncRNAs may be more likely than in intergenic regions.

To conclude, bifunctional (moonlighting) lncRNAs indeed exist. Identifying them, analysing the properties of the proteins they encode, and understanding their functional regulation is an important avenue for future research.

## Methods

### Mathematical model

I calculated the steady state solutions to the mathematical model by analytically solving the ordinary differential equations (Equations 1 – 5).

### Preparation of the mRNA and ncRNA sequence data

I downloaded *D. melanogaster* mRNA and ncRNA sequences directly from FlyBase (release dmel-6.46, Gramates *et al*., 2022). To obtain human mRNAs and lncRNA sequences, I downloaded the gene annotation (gtf) and chromosome sequences from the RefSeq database (release GRChg38.p14, O’Leary *et al*., 2015). Next, I extracted mRNAs co-ordinates from the gtf file, based on their transcript accession number starting with “NM”. Similarly, I extracted the co-ordinates of non-coding RNAs (ncRNAs) based on their transcript accession number starting with “NR”. I excluded the annotated pseudogenes that also have an “NR” accession number. Next, I extracted mRNA and ncRNA sequences using the program *gff-read* (version 0.12.7 Pertea and Pertea, 2020). I further extracted lncRNA sequences that are ncRNAs longer than 200nt. I obtained functional human lncRNAs from the lncRNAWiki2.0 database (Liu *et al*., 2021). Specifically, I extracted the lncRNAs gene names, that were annotated to interact with a protein coding gene, an RNA, or a protein. Next, I downloaded the transcripts arising from these genes from the Ensembl database (Harrison *et al*., 2023) using BioMart web interface.

I extracted ORF sequences from each of these datasets using the program *getorf* and translated them into protein sequences using *transeq*. Both these programs are a part of the EMBOSS suite (version 6.6.0.0, Rice *et al*., 2000). I estimated the strength of Kozak consensus sequences (KCS) using the data from Acevedo *et al*. (2018, *D. melanogaster*) and Noderer *et al*. (2014, *H. sapiens*).

### Estimating the properties of lncRNA encoded proteins

#### Hydropathy index

Hydropathy index for each amino acid had been estimated by their partitioning between water and octanol (Wimley *et al*., 1996). Using this scale, I estimated the average hydropathy index for a protein sequence from each of its constituent amino acid.

#### Isoelectic point

I estimated isoelectic point using the program *pepstats* from the EMBOSS suite (version 6.6.0.0, Rice *et al*., 2000).

#### Shannon entropy

I estimated Shannon entropy of a protein sequence using the frequency of every “word” ranging from 1 to 5 amino acids. Specifically, if *W*_*i*_ denotes all the unique words of length *i* in a protein sequence and 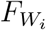 denotes their frequency, then I calculated Shannon entropy *H*, as:

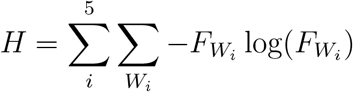

#### Linguistic complexity

If *W*_*i*_ denotes all the unique words of length *i* in a protein sequence of length *k*, 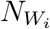 denotes their counts, and *U* (*i, k*) denotes the maximum number of possible words of length *i* in the protein sequence, then I calculated linguistic complexity *L* as:

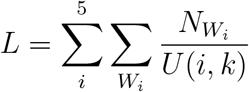

The total number of words of length *i* in a sequence of length *k*, is *k − i* + 1. The total number of possible protein words of length *i* is 20^*i*^. *U* (*i, k*) is the smaller value between *k − i* + 1 and 20^*i*^.

#### Structural propensity/disorder

I used the machine learning based software tool ADOPT to predict a protein’s structural disorder from its sequence (Redl *et al*., 2023). This tool extracts amino acid level sequence representations from the ESM-1 protein language model, and uses nuclear magnetic resonance (NMR) data sets to train its model. For every amino acid in a protein sequence, ADOPT outputs a score. A score smaller than 8 indicates that the amino acid is present in a disordered region, and *vice versa*. For every lncRNA encoded protein sequence, I estimated the median value of the ADOPT scores from its constituent amino acids. Because larger score values indicate the likelihood of a structure, I refer to the median ADOPT score for a protein as its “structural propensity”.

#### Nearest protein domains in the sequence space

First, I downloaded the sequences of the ECOD domains, and narrowed the list of sequences to those that are 10 – 120 amino acids long. Next, I used ESM2 language model to generate embeddings for the ECOD domains, as well as for the lncRNA encoded protein sequences. Next, I calculated the Manhattan distance between the embeddings of an lncRNA encoded protein and that of the different ECOD domains, and thus identified the nearest domain. A Manhattan distance (L1 norm) is a better distance metric for multidimensional data than Euclidean distance (L2 norm) because the latter is more sensitive to outliers (Barrodale, 1968).

#### Structure predictions

I used the ESMFold web interface for the structure predictions.

## Code and data availability

All data and used in this study are already published and are publicly available. Programming scripts used for analysis of hydropathy, Shannon entropy, and linguistic complexity can be found in the public github repository *BharatRaviIyengar/useful awks* (*hydropathy*.*awk, complexity*.*awk*). Other analyses have been performed using publicly available software.

